# Informational and Normative Influence on Conformity in Autism

**DOI:** 10.64898/2025.11.30.691476

**Authors:** Akiko Kobayashi, Takuya Makino, Yuka Mizuno, Hirotaka Kosaka, Keise Izuma

## Abstract

This preregistered study examined whether adults with autism spectrum disorder (ASD) show reduced social conformity and whether any such reduction depends on the type of social influence. Social conformity, defined as the tendency to adjust judgments to align with those of others, is typically driven by normative (acceptance-seeking) and informational (accuracy-seeking) motives. Thirty adults with ASD and 30 matched neurotypical (NT) adults completed two tasks: a preference rating task indexing normative influence, and a dot-counting task indexing informational influence with monetary rewards. Contrary to our predictions, adults with ASD conformed as much as NT adults in the preference rating task but showed significantly reduced conformity in the dot-counting task. Exploratory analyses indicated that this reduction was driven by a distinct subgroup of nine adults with ASD who never revised their initial estimates despite informative social cues, resulting in poorer accuracy and lower rewards. When this subgroup was excluded, group differences in conformity were no longer evident.

These findings suggest that, overall, adults with ASD are as susceptible as NT adults to normative influence but less responsive to informational influence, highlighting the importance of distinguishing between types of social influence and considering individual differences when characterizing social behavior in ASD.

## Introduction

Our decisions and behaviors are often shaped by others’ judgments, opinions, beliefs, and actions. When we encounter views that differ from our own—even when they are incorrect—we frequently reconsider and adjust our perspectives to align with others, a phenomenon known as social conformity (Cialdini & Goldstein, 2004).

Conforming to others is a fundamental form of social learning that supports complex social functioning (Shamay-Tsoory et al., 2019). In contemporary contexts, conformity serves multiple functional purposes: social conformity plays a critical role in promoting group cohesion (Lott & Lott, 1961), enhancing information processing—particularly under conditions of uncertainty or when individual learning is biased (Toelch et al., 2014; Toyokawa & Gaissmaier, 2022), and promoting cooperative behavior (Parks et al., 2001) and prosocial behavior (Xiao, 2017). Conversely, a diminished tendency to conform may pose certain risks, including difficulties in fitting within a group and the potential for ridicule or negative evaluation by group members (Deutsch & Gerard, 1955; Shamay-Tsoory et al., 2019; Haihambo et al., 2025).

Individuals with autism spectrum disorder (ASD) are considered to exhibit a reduced tendency to conform. ASD is a neurodevelopmental disorder characterized by deficits in social communication and interaction, as well as restricted and repetitive behaviors. Individuals with ASD possess certain characteristics that may lead to a reduced tendency to conform to others. For example, they tend to pay less attention to social cues (Hedger et al., 2020) and are less responsive to opportunities for social approval (Izuma et al., 2011). According to the social motivation account of ASD (Chevallier et al., 2012), these characteristics reflect reduced social orienting and seeking, stemming from diminished social motivation—such as a reduced desire to gain acceptance or avoid rejection. Using structured teacher interviews, Nenniger & Müller (2021) reported that students with ASD are generally less susceptible to peer influence. To the extent that conformity is driven by the motivation to enhance social acceptance, individuals with ASD would be expected to exhibit reduced conformity due to their diminished social motivation.

Understanding this reduced tendency to conform is important not only for clarifying the mechanisms underlying social adaptation difficulties in ASD but also for informing the development of targeted interventions and support strategies aimed at enhancing social participation and well-being. However, empirical research specifically examining conformity in ASD is relatively limited, and existing findings have been mixed. Bowler & Worley (1994) examined social conformity in adults diagnosed with Asperger syndrome using a version of the classic Asch (1952) paradigm, in which participants performed a simple line judgment task while, on certain trials, four confederates intentionally gave incorrect answers before the participant responded.

Although the study did not find a statistically significant difference between the Asperger and control groups—likely due to the small sample size (eight adults with Asperger syndrome)—the pattern of results was suggestive: all 17 control participants conformed at least once to the incorrect judgments, whereas only half of the participants with Asperger syndrome did so. Yafai et al. (2014) conducted a similar study with children (n=15 per group) and found a significantly reduced tendency to conform in the ASD group compared with controls.

Lazzaro et al. (2019) investigated conformity tendencies in adults with ASD using a memory-based conformity task and found comparable levels of conformity between the ASD and control groups. Participants were first exposed to a list of words and then completed a recognition memory test while viewing the responses of four other individuals. A computer-generated condition was also included, in which responses were presented as coming from four computer programs. The study found that both groups conformed more to human opinions than to computer-generated ones, with no significant difference between the ASD and control participants.

One previously underexplored factor that may contribute to inconsistencies in the literature on conformity in ASD is the presence of two distinct motivations underlying conformity. Social conformity is thought to be driven by two separate influences: informational and normative (Deutsch & Gerard, 1955). Informational influence refers to the motivation to be accurate, while normative influence reflects the motivation to be accepted by others. In the classic Asch line judgment task, as used by Bowler and Worley (1994) and Yafai et al. (2014), the task is easy and therefore conformity is typically interpreted as reflecting normative influence. By contrast, in more difficult tasks—such as Lazzaro et al.’s (2019) memory test, in which participants’ baseline accuracy was below 70% (with a chance level of 50%)—conformity is more likely driven by informational influence.

According to the social motivation theory of ASD (Chevallier et al., 2012), reduced conformity in ASD may emerge primarily in situations where normative influence plays a dominant role, as such situations rely heavily on the motivation to enhance social acceptance. By contrast, conformity driven by informational influence—where individuals rely on others’ input to improve task performance—may not differ substantially between individuals with ASD and neurotypical (NT) controls. In the present study, we tested this hypothesis in adults with ASD using two experimental tasks designed to dissociate these influences: a preference rating task and a dot-counting task (Figure 1).

**Figure 1.**
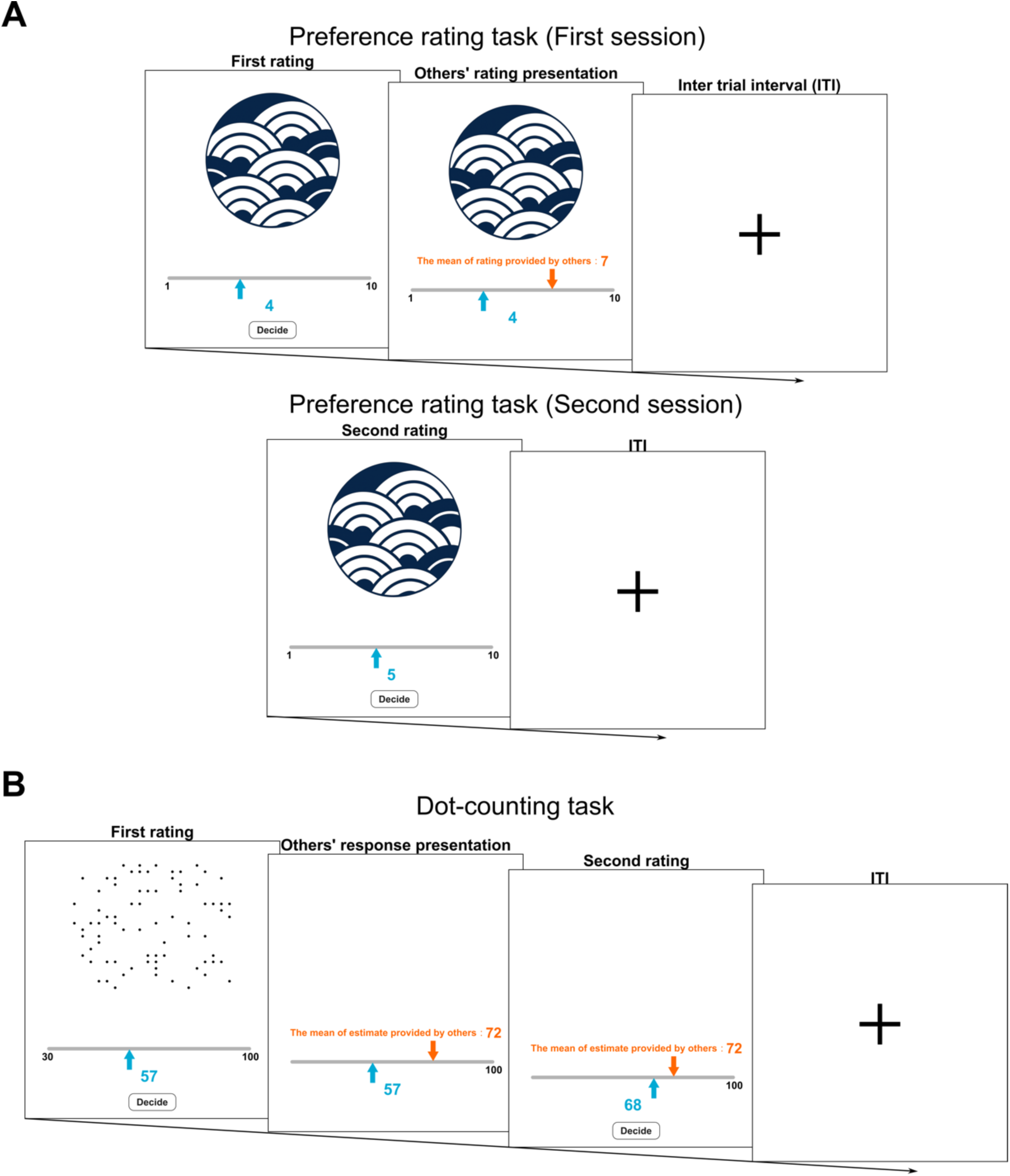
Images of the preference rating and dot-counting task (A) An example of the preference rating task. During the first session of the preference rating task, participants were requested to assess their level of preference for each design by moving the blue arrow along a scale ranging from 1 (Strongly Dislike) to 10 (Strongly Like). Immediately after the submitting the initial rating, a participant viewed the rating provided by other individuals for the same T-shirt, indicated by the orange arrow. The blue number indicates the rating selected by the participant. In the second session, participants were requested to assess their level of preference for the same design presented in the first session, without being shown others’ rating. (B) An example of the dot-counting task. During the dot-counting task, a participant was prompted to estimate the quantity of dots displayed on the screen by moving the blue arrow along a scale ranging from 30 to 100. Immediately after the submitting the initial rating, the participant viewed the rating provided by other individuals for the same dot image, indicated by the orange arrow. The blue number indicates the number selected by the participant. Participants were given an opportunity to revise their first rating after viewing the ratings provided by others.

In the preference rating task (Izuma & Adolphs, 2013), participants were shown various T-shirt designs and asked to rate how much they liked each one. After providing their initial ratings, they were shown the ratings provided by others (i.e., group norm). Later, in a separate session, they had an opportunity to re-rate the same T-shirts. Fashion preferences, like other hedonic or aesthetic judgments, are especially susceptible to normative social influence because these judgments are largely based on subjective feelings rather than objective criteria (Venkatesan, 1966). While informational influence can play a significant role in decisions involving utilitarian products—where others’ opinions can provide useful information for making objectively better choices (e.g., ratings on an online shopping site for electronic products)—it is generally weaker for purely subjective, hedonic evaluations such as clothing preference. Furthermore, similar subjective preference-rating tasks have been widely used in functional magnetic resonance imaging (fMRI) studies of conformity (Berns et al., 2005; Campbell-Meiklejohn et al., 2010; Izuma & Adolphs, 2013; Klucharev et al., 2009; Levorsen et al., 2021; Nook & Zaki, 2015). These studies have consistently demonstrated robust social conformity in domains involving subjective preferences (e.g., food, aesthetics, fashion), even when other raters were not physically present. A meta-analysis of neuroimaging studies on social conformity by Wu et al. (2016) revealed that agreement between one’s own rating and the group’s rating activated the ventral striatum—a core region of the brain’s reward network—suggesting that alignment with group norms is experienced as rewarding, possibly reflecting anticipated social approval or relief at avoiding deviation from the norm. Thus, normative influence appears capable of operating even in minimal or non-interactive contexts, driven by internalized motivations to align with group norms or avoid perceived deviance. Given these considerations, conformity in the preference rating task is assumed to primarily reflect normative influence. Owing to the diminished social motivation observed in individuals with ASD, we hypothesized that they would show reduced conformity in the preference rating task compared with NT individuals.

By contrast, in the dot-counting task (Fliessbach et al., 2007), participants viewed briefly presented stimuli (500 ms each), each containing up to 90 dots, and estimated the total number. After providing their estimate, participants were presented with the estimates of other participants and were given the opportunity to revise their own.

Because this task has objectively correct answers, conformity observed here is assumed to primarily reflect informational influence. To keep the task neither trivially easy nor excessively difficult, the exposure duration and numerosity range were chosen to yield a moderate difficulty level, minimizing floor/ceiling effects. To further enhance participants’ motivation for accuracy, monetary incentives were provided based on the proximity of their estimates to the correct value. Although some neuroimaging studies have found differences in neural responses to monetary rewards between ASD and NT individuals (see Clements et al., 2018 for a meta-analysis), these studies reported similar incentive-related performance improvements (e.g., faster reaction times) across these groups (Delmonte et al., 2012; Dichter, Felder, et al., 2012; Dichter, Richey, et al., 2012; Scott-Van Zeeland et al., 2010). Therefore, we predicted comparable conformity levels between individuals with ASD and NT controls in the dot-counting task.

To summarize, our preregistered predictions were as follows: individuals with ASD would show reduced conformity in the preference rating task but comparable conformity to NT controls in the dot-counting task.

## Materials and Methods

All procedures were conducted in accordance with the Declaration of Helsinki (2013), and ethical approval for the study was granted by ethics committees of Kochi University of Technology and Fukui University. Participants provided written informed consent before their engagement.

### Preregistration

We preregistered the hypotheses, study design, data collection procedures, sample size, analysis plan, and participant exclusion criteria on the Open Science Framework platform (https://doi.org/10.17605/OSF.IO/GXKBW). Unless otherwise noted, we analyzed the data in accord with the preregistration.

### Participants

As preregistered, we recruited 30 (24 male) participants who were diagnosed with ASD based on the Diagnostic and Statistical Manual of Mental Disorders, 5th Edition (DSM-5) (APA, 2013). Their diagnosis was confirmed using the Diagnostic Interview for Social and Communication Disorders (DISCO; Wing et al., 2002) by a psychiatrist (H.K.). We also recruited 31 (23 male) NT adults. Sample size was determined based on previous studies of social conformity in ASD (Large et al., 2019; Lazzaro et al., 2019). Intellectual functioning was assessed with standardized Japanese versions of the Wechsler Adult Intelligence Scales (WAIS): either a short form of the WAIS-III (Fujita et al., 2011) or the full WAIS-IV (Wechsler, 2008), depending on when participants were tested. We confirmed that all participants had a full-scale IQ of at least 70, indicating the absence of intellectual disability or other psychiatric disorder. One NT participant was excluded from analyses based on a preregistered exclusion criterion (see “Exclusion criteria” below for more detail). Our final sample comprised 30 ASD and 30 NT participants. Two groups were matched on age, gender ratio, and full-scale IQ (see Table 1).

**Table 1.** Participant demographics.

|  | ASD (n=30) |  | NT (n=30) |  | *T* | *P* |
| --- | --- | --- | --- | --- | --- | --- |
|  | Mean | SD | Mean | SD |  |  |
| Age | 33.43 | 4.64 | 31.93 | 7.15 | 0.96 | 0.339 |
| FSIQ | 108.43 | 12.07 | 110.43 | 13.17 | −0.61 | 0.542 |

*Note.* FSIQ denotes full-scale IQ. FSIQ was measured by the WAIS-III or WAIS-IV. ASD: Autism Spectrum Disorder, NT: neurotypical controls, *T*: t-value, *P*: p-value and SD: standard deviation.

### Task procedure

We employed two types of conformity tasks: the preference rating and dot-counting tasks (Figure 1). In the preference rating task (Figure 1A), we utilized the social conformity task employed in our previous study (Izuma & Adolphs, 2013). The preference rating task consisted of two sessions, each with 100 trials. In the first session, participants were presented with a front image of a T-shirt, accompanied by a 10-point scale. Participants were instructed to evaluate their preference for each image using the scale that ranges from 1 (Strongly Dislike) to 10 (Strongly Like). Participants were required to provide their response by adjusting the slider, and subsequently clicking the “Decide” button to submit their rating, with no time constraint imposed. Immediately after the submission of the initial rating, the rating provided by other individuals for the same T-shirt was displayed for a duration of 1,700 ms. There was an inter-trial interval (ITI) for 500 ms between trials. Participants were informed that an orange cursor in each trial symbolizes “the aggregated opinion of individuals who previously took part in the same experiment.” In reality, it was systematically manipulated based on a similar method to that employed in our previous study (Levorsen et al., 2021). The ratings provided by the group aligned exactly with each participant’s own rating in approximately 17.5%–25% of trials. In the remaining 75%–82.5% of trials, the group ratings were distributed approximately evenly above or below the participant’s rating, with differences never exceeding three points. After a brief break, participants took part in the second session where they were unexpectedly asked (without prior announcement during the initial instructions) to reevaluate each of the 100 images using the same 10-point scale. However, this time, others’ ratings were not presented.

In the dot-counting task (Figure 1B), we utilized a modified version of the estimation task used in the previous study (Fliessbach et al., 2007). The dot count task consists of 100 trials. Participants were presented a series of black dots for a duration of 500 ms. Following stimulus presentation, participants were prompted to estimate the quantity of dots displayed on the screen. Participants were required to select a numerical value within the range of 30–100 by manipulating a slider with a blue cursor, followed by clicking the “Decide” button to submit their initial rating, with no imposed time constraint. Following the submission, the average estimate ostensibly provided by other individuals, indicated by an orange cursor, was displayed. As in the preference rating task, participants were informed that “the other individuals” were those who had participated in the same experiment. In reality, others’ ratings were chosen at random within a range defined by the actual dot count ± 20, following a uniform distribution.

Subsequently, participants were given the opportunity to revise their initial response. Irrespective of whether participants modified their response, they had to click the “Decide” button again to submit their final answer, with no imposed time constraint. No trial-by-trial feedback about correctness (e.g., the true dot count) was provided at any point, preventing learning across trials. There was an ITI for 500 ms between trials. We prepared two distinct stimulus sets, each containing 50 unique images depicting between 41 and 90 black dots, without overlap between the two sets (totaling 100 images). To ensure task difficulty, no images contained fewer than 41 dots. During the first half of the task, participants viewed each image once, with images randomly selected from one stimulus set. After a brief break, the second half of the task proceeded similarly using the other stimulus set. Participants were informed beforehand that they could earn monetary bonus based on the accuracy of their final estimates, and were encouraged to respond as accurately as possible, although specific criteria for earning bonuses were not disclosed. In reality, participants earned 1 point when their final estimate fell within ±5 dots of the actual number, and each point was converted to a monetary bonus of 10 yen. For both tasks, stimuli were presented using jsPsych.

### Exclusion criteria

We calculated the correlation coefficient of the first and second ratings for the preference rating task (i.e., *r_PreferenceRating_*) and the correlation coefficient of the first response and actual number of dots for the dot-counting task (i.e., *r_DotCounting_*). Based on the results of a pilot study, we decided to exclude participants who exhibited an *r_PreferenceRating_* lower than 0.35 or an *r_DotCounting_* lower than 0.42 from the main analysis. Data from one participant from the NT group were excluded from the main analysis because s/he met the preregistered exclusion criteria for the dot-counting task.

## Statistical analysis

As preregistered, we calculated the conformity tendencies using multiple regression analysis for each task, performed individually for every participant, as follows:

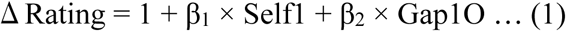

where *ΔRating* represents the difference between the participant’s first and subsequent ratings (second rating – first rating), *Self1* denotes the participant’s first rating, and *Gap1O* denotes the difference between the participant’s first rating and the ratings from other individuals (other people’s rating – participant’s first rating). To verify that conformity tendencies measured by the two tasks—each designed to assess normative and informational influences, respectively—did not reflect a common underlying tendency, we conducted a correlation analysis between conformity tendencies (i.e., β_2_) obtained from each task.

In our preregistration, we planned to quantify conformity tendencies using multiple regression analyses performed separately for each participant and task, followed by a group-level comparison implemented through a mixed two-way analysis of variance (ANOVA), with group (ASD and NT) as a between-subject factor and task type (preference rating and dot-counting tasks) as a within-subject factor. However, given that generalized linear mixed models (GLMMs) provide greater statistical power than traditional summary-based approaches—such as ANOVAs conducted on per-subject means—by modeling trial-level data and accounting simultaneously for random effects due to subjects and items (Barr et al. 2013; Baayen et al., 2008), we instead opted to employ GLMMs as follows (for details of the preregistered analyses and corresponding results, see the Supplemental Information):

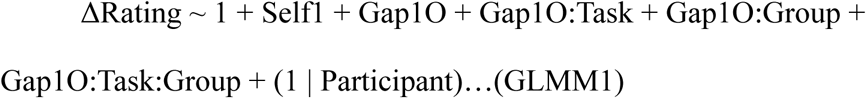

where *ΔRating* represents the difference between the participant’s first and subsequent ratings (second rating – first rating), *Self1* denotes the participant’s first rating (i.e., controlling for a regression-to-the-mean effect), and *Gap1O* denotes the difference between the participant’s first rating and the ratings from other individuals (other people’s rating – participant’s first rating). *Task* and *Group* are dummy variables indicating the type of tasks (Preference rating task coded as 1; Dot-counting task coded as 0) and whether the participant has been diagnosed with ASD (ASD group coded as 1; NT group coded as 0). The interaction between *Gap1O*, *Task*, and *Group* in the GLMM1 corresponds to a mixed two-way ANOVA interaction effect, as it captures how the relationship between *Gap1O* (i.e., predictor of conformity) and the outcome variable varies across the levels of *Task* (within-subject factor) and *Group* (between-subject factor). All predictors, including *ΔRating*, *Self1*, and *Gap1O*, were standardized prior to analysis.

We applied the conventional significance threshold *p* <.05 for GLMM and post-hoc tests. To control for the type 1 error rate in our analyses, we implemented the Bonferroni correction.

### Additional analysis (not preregistered)

We employed two different tasks to examine group differences in conformity tendencies driven by two distinct types of influences. To assess whether the conformity behavior observed in each task was driven by distinct influences, we first conducted correlation analyses between conformity tendencies (β_2_) obtained from each task.

In the dot-counting task, participants received an additional bonus based on their estimation accuracy. We conducted a group comparison of the additional bonus by performing a two-sample t-test. Further, to assess whether adjusting participants’ initial responses based on others’ responses contributed to greater accuracy, we conducted correlation analyses between conformity tendencies in the dot-counting task and deviations of the second responses from the actual number of presented dots within each group.

### Results Group comparison of conformity tendency

To examine whether individuals with ASD exhibit distinct patterns of conformity depending on the task, we constructed GLMM1, which first revealed a strong positive main effect of Gap1O (*b* = 0.46, *t* = 28.36, *p* <.001; Table 2): larger discrepancies between participants’ initial responses and the group’s responses predicted greater conformity. The main effect of *Self1* was also significant (*b* = −0.17, *t* = −20.32, *p*

**Table 2.** Results of GLMM1 examining group differences in conformity tendencies between the preference rating and dot-counting tasks.

|  | Estimate | SE | *t* | *p* | 95% CI |
| --- | --- | --- | --- | --- | --- |
| Intercept | 0.00 | 0.01 | 0.00 | 1.000 | [−0.02 0.02] |
| Self1 | −0.17 | 0.01 | −20.32 | < .001 | [−0.19 −0.15] |
| Gap1O | 0.46 | 0.02 | 28.36 | < .001 | [0.43 0.49] |
| Gap1O:Task | −0.33 | 0.02 | −14.33 | < .001 | [−0.37 −0.28] |
| Gap1O:Group | −0.16 | 0.02 | −6.98 | < .001 | [−0.21 −0.12] |
| Gap1O:Task:Group | 0.14 | 0.03 | 4.42 | < .001 | [0.09 0.21] |

*Note.* Self1: first rating, Gap1O: difference between first rating and others' rating, Task: dummy code of task (preference rating:1; dot-counting:0), Group: dummy code of group (ASD:1; NT:0), SE: standard error, CI: confidence interval.

<.001), indicating a significant regression-to-the-mean effect, consistent with our previous studies (Izuma & Adolphs, 2013; Levorsen et al., 2021). The two-way interaction between *Gap1O* and *Task* was significant (b = −0.33, *t* = −14.33, *p* <.001), indicating that the strength of the conformity effect varied across the tasks. The two-way interaction between *Gap1O* and *Group* was also significant (*b* = −0.16, *t* = −6.98, *p* <.001), indicating that the strength of the conformity effect varied across groups. Importantly, the significant two-way interactions were qualified by a significant three-way interaction between *Gap1O*, *Task*, and *Group* (*b* = 0.14, *t* = 4.42, *p* <.001; Table 2). This significant three-way interaction indicates that the effect of *Gap1O* (i.e., the discrepancy between participants’ initial responses and the group’s responses) on conformity differs across tasks and that this task-dependent effect further varies between groups. In other words, the pattern of conformity across groups is modulated by task context (Figure 2).

**Figure 2.**
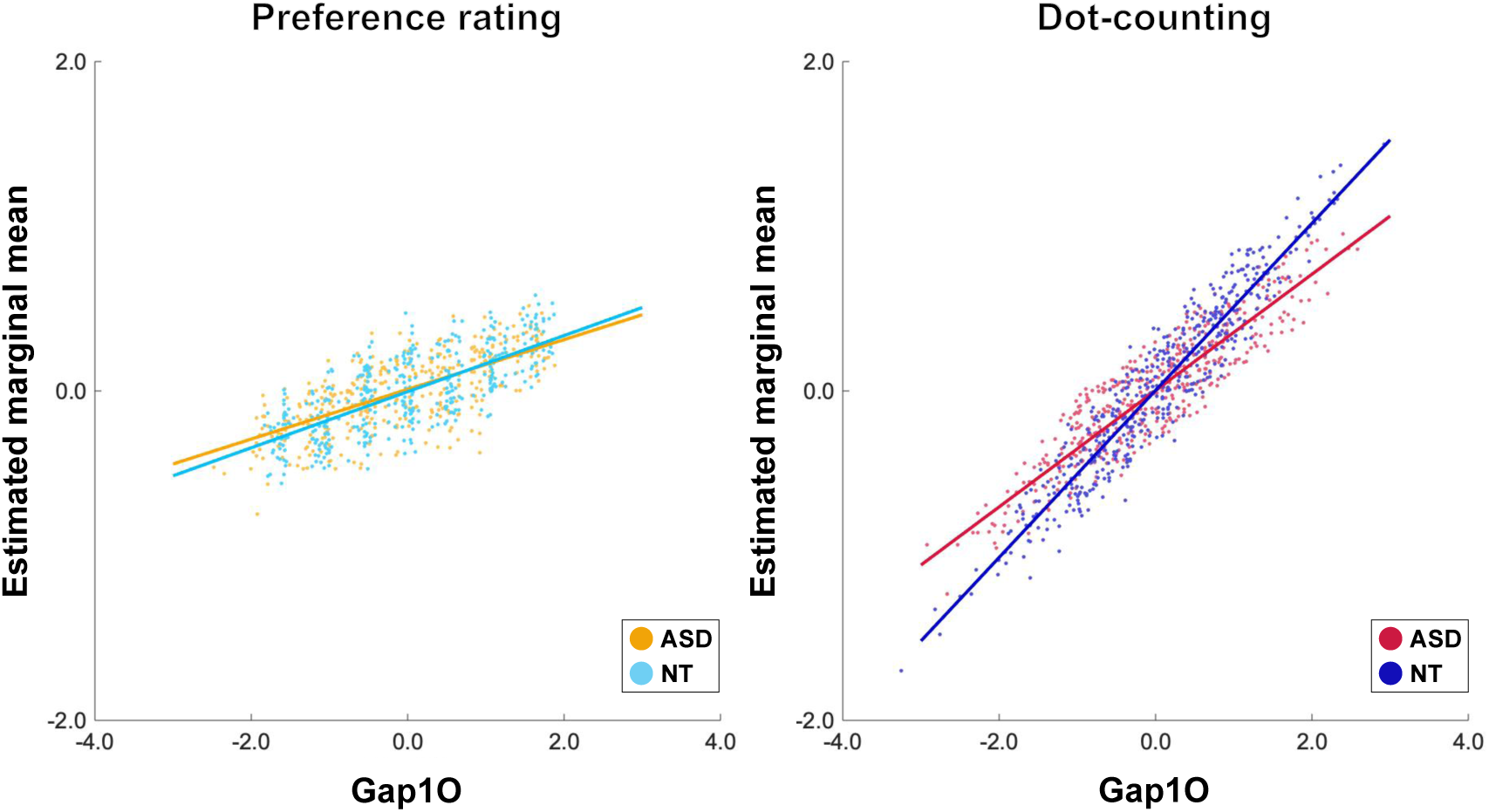
Estimated marginal means of rating update calculated from GLMM1, plotted by group and task Estimated marginal means of rating update are calculated from GLMM1. Each plot represents a data corresponding to an individual trial. The x-axis represents the standardized difference between others’ rating and the first rating (Gap1O), while the y-axis indicates estimated marginal means of rating update. 500 data points were randomly sampled from each group and task (from 3000 per condition) to illustrate the interaction.

To examine the three-way interaction effect in greater detail, we additionally constructed separate GLMMs for each task (GLMM1_PreferenceRating_ and GLMM1_DotCounting_). The model specification for both was as follows (all terms are defined as in GLMM1):

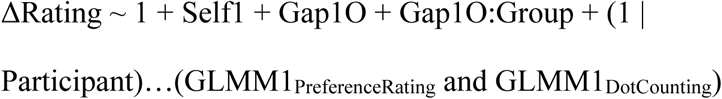

The result first revealed no significant interaction between *Gap1O* and *Group* in the preference rating task (*b* =-0.02, *t* = −0.87, *p_adjusted_* =.764; see Table 3), indicating no significant group difference in conformity tendency in the preference rating task. By contrast, there was a significant interaction between *Gap1O* and *Group* in the dot-counting task (*b* = −0.16, *t* = −7.92, *p_adjusted_* <.001; see Table 4), indicating that individuals with ASD exhibited a significantly weaker conformity tendency than NT individuals in the dot-counting task.

**Table 3.** Results of GLMM1_PreferenceRating_ testing group differences in conformity tendencies in the preference rating task.

|  | Estimate | SE | *t* | *p*<sub>adjusted</sub> | 95% CI |
| --- | --- | --- | --- | --- | --- |
| Intercept | 0.00 | 0.01 | 0.00 | 1.000 | [−0.02 0.02] |
| Self1 | −0.31 | 0.01 | −25.24 | < .001 | [−0.34 −0.29] |
| Gap1O | 0.11 | 0.02 | 6.31 | < .001 | [0.08 0.14] |
| Gap1O:Group | −0.02 | 0.02 | −0.87 | .764 | [−0.07 0.03] |

*Note.* Self1: first rating, Gap1O: difference between first rating and others' rating, Group: dummy code of group (ASD:1; NT:0), SE: standard error, CI: confidence interval.

**Table 4.**
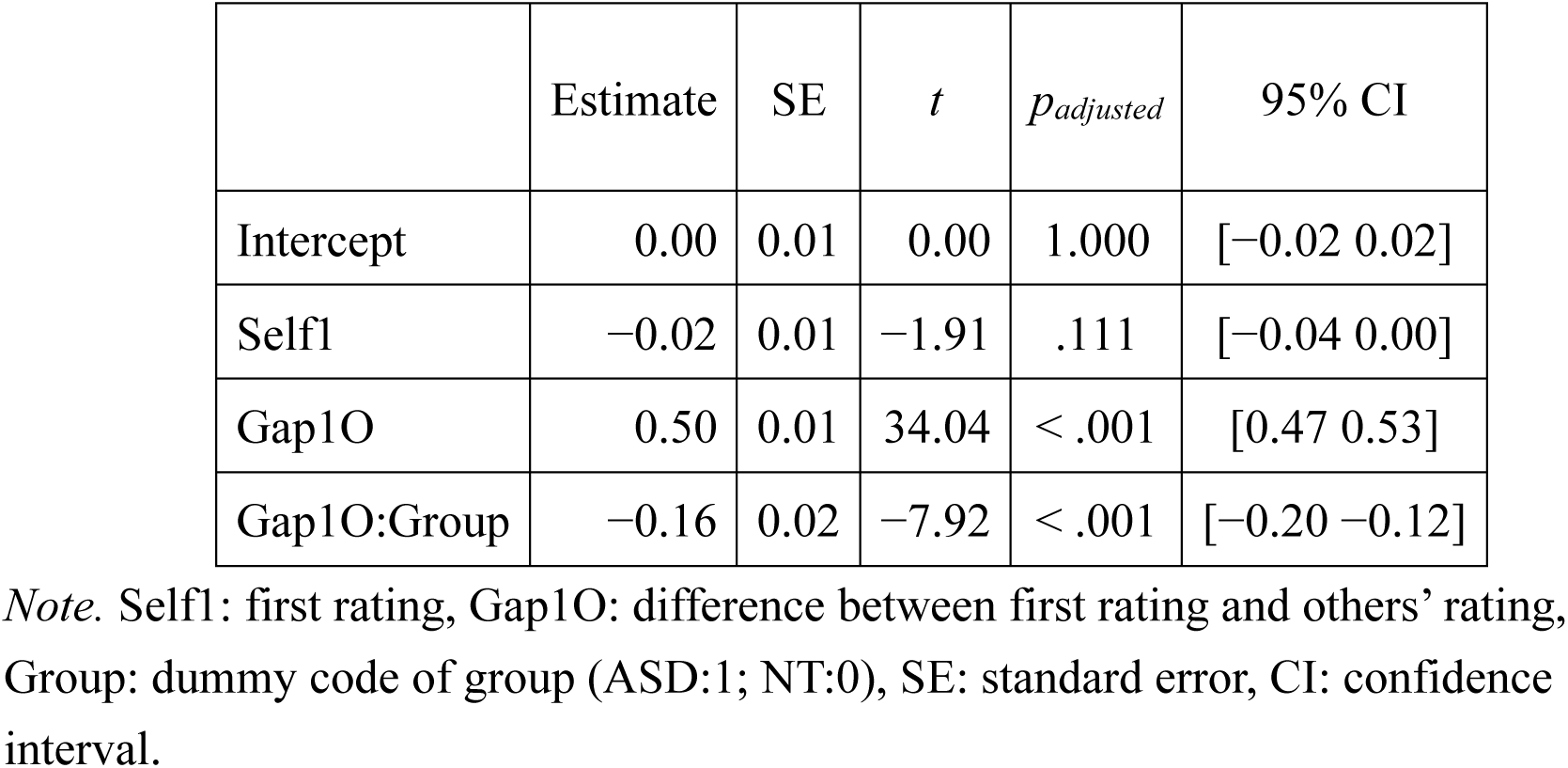
Results of GLMM1_DotCounting_ testing group differences in conformity tendencies in the dot-counting task.

Taken together, these results were in the opposite direction of our preregistered predictions: rather than showing reduced conformity in the preference rating task and comparable conformity in the dot-counting task, individuals with ASD exhibited no significant reduction in conformity in the preference rating task but showed markedly weaker conformity in the dot-counting task.

## Exploratory analyses (not preregistered)

### Correlation between conformity tendencies in the preference rating and dot-counting tasks

To examine whether conformity behavior observed in each task was driven by distinct motivations (e.g., accuracy vs. social acceptance), we conducted correlation analyses between conformity tendencies (β₂, calculated from Equation 1) obtained from the preference rating and dot-counting tasks. Neither the ASD nor the NT group showed a significant correlation between conformity tendencies across tasks (ASD: *r* =.26, *p* =.160; NT: *r* =.18, *p* =.537; Figure 3A), suggesting that conformity in each task may be guided by different underlying motivations. We confirmed that these results were replicated even when excluding those who never modified their initial ratings during the dot-counting task (ASD: *r* =.22, *p* =.342; NT: *r* =.31, *p* =.105).

**Figure 3.**
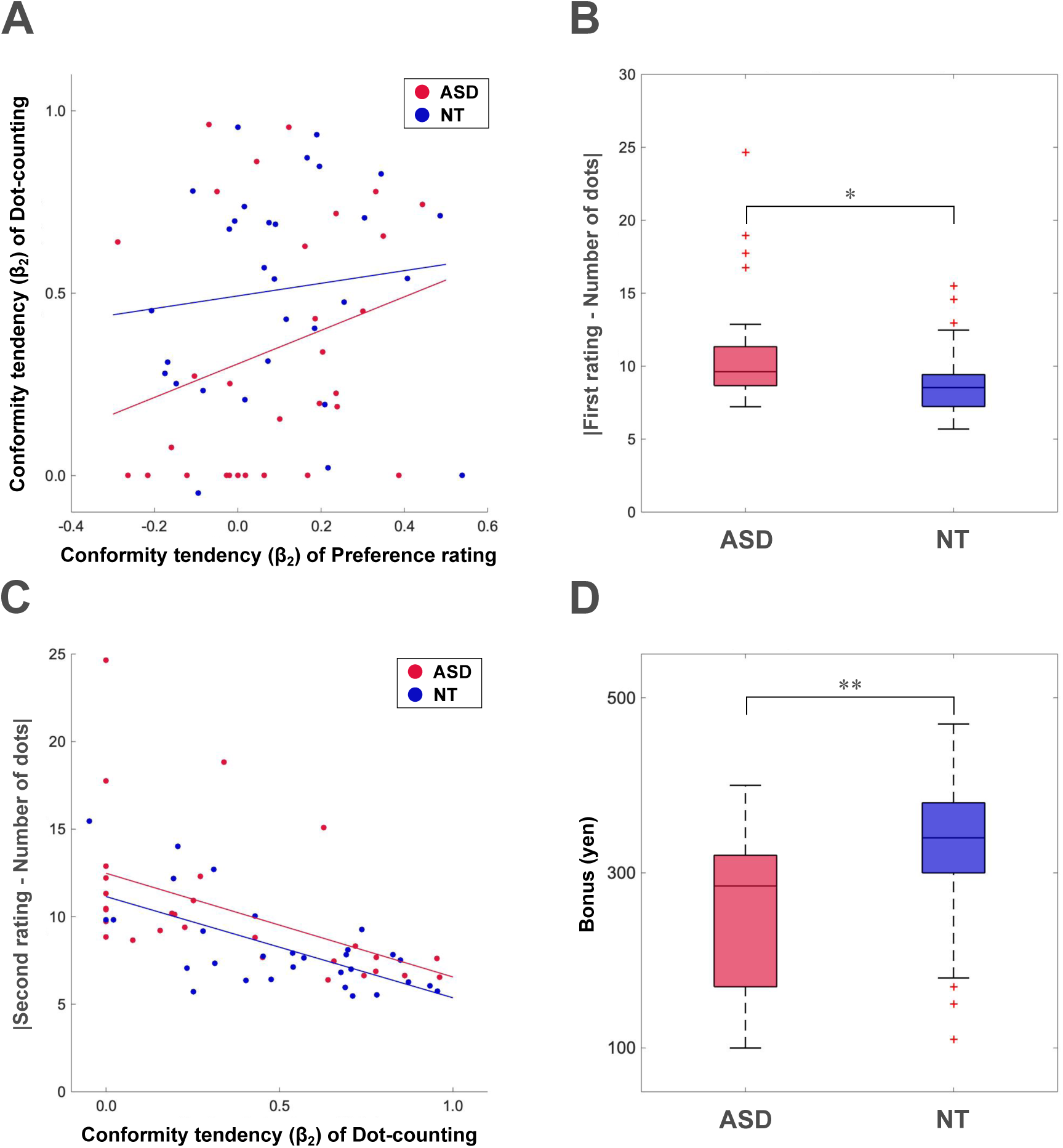
Results of exploratory analyses on task performance (A) Relationship between conformity tendencies of each task. The x-axis represents the conformity tendencies obtained from the preference rating task, while the y-axis represents the conformity tendencies obtained from the dot-counting task. Conformity tendencies of both tasks were derived from Equation 1 (β₂). Each plot represents the data of an individual participant. (B) Results of group comparison on initial estimation accuracy in the dot-counting task (i.e., deviations of initial responses from the actual number of presented dots). (C) Relationship between conformity tendencies in the dot-counting task and second estimation accuracy in the dot-counting task. The x-axis represents the conformity tendencies obtained from the dot-counting task, while the y-axis represents deviations of second responses from the actual number of presented dots. Each plot represents the data of an individual participant. (D) Results of group comparison on the additional bonus earned based on estimation accuracy in the dot-counting task. The box indicates the interquartile range (25th–75th percentile), and the central line represents the median. Whiskers extend to 1.5 × IQR, and red crosses indicate outliers. ^∗^*p* <.05, ^∗∗^*p* <.01.

### Were individuals with ASD better at estimating the number of dots in the dot-counting task?

One possible reason for the findings that individuals with ASD exhibited a reduced tendency to conform in the dot-counting task may be because they were better at the task, so they did not have to rely on others’ opinion. To examine this possibility, we compared estimation errors in the dot-counting task between groups, defined as the absolute discrepancy between the first rating and the actual number of dots. The result showed that estimation errors of initial ratings were significantly larger for individuals with ASD compared with NT individuals (*t* (58) = 2.62, *p* =.014; Figure 3B), indicating that the former were actually worse at the dot-counting task. Because others’ ratings were determined based on the actual number of presented dots, we confirmed that the discrepancy between participants’ initial ratings and others’ ratings was marginally larger in individuals with ASD compared with NT individuals (*t* (58) = 1.93, *p* =.058), indicating that individuals with ASD perceived larger deviations between their own and others’ opinions. Taken together, these findings indicate that the reduced conformity among individuals with ASD in the dot-counting task cannot be attributed to superior task performance, and in fact, reduced conformity occurred despite their larger estimation errors and greater disagreement with group ratings.

Notably, in both groups, conformity tendency was significantly correlated with task performance: the greater the conformity to group estimates, the smaller the estimation error (ASD: *r* = −.49, *p* =.006; NT: *r* = −.65, *p* <.001; Figure 3C). Thus, as a result of both poorer initial estimation accuracy and lower conformity, individuals with ASD earned fewer points (i.e., less monetary bonus) in the dot-counting task compared with NT individuals (*t* (58) = −3.01, *p* =.003; Figure 3D).

### Does a subgroup of individuals with ASD explain the reduced conformity in the dot-counting task?

One important difference between the two tasks was that the dot-counting task allowed participants to immediately modify their rating after viewing others’ ratings (Figure 1), making conformity much more explicit in the dot-counting task than in the preference rating task. Visual inspection of the data revealed that 9 out of 30 individuals with ASD (compared with only 1 out of 30 NT individuals) never modified their initial ratings during the dot-counting task, indicating a complete absence of conformity in these participants. Those individuals who showed a strong consistency bias completely ignored others’ ratings, even though these ratings were informative and could have helped them earn a greater monetary bonus. A chi-squared test comparing the proportion of individuals who never revised their rating confirmed a significant group difference (χ*²*(60) = 7.68, *p* =.006). For clarity, we refer to these individuals as the ASD-never-conforming subgroup, and to the remaining 21 ASD group participants who revised at least one of their ratings as the ASD-conforming subgroup.

We further examined the distribution of conformity tendencies in the dot-counting task separately for individuals with ASD and NT individuals. The conformity scores of NT individuals did not significantly deviate from normality (Shapiro–Wilk test, *W* =.97, *p* =.250), whereas those of individuals with ASD showed a clear deviation from normal distribution (*W* =.86, *p* <.001; Figure 4B). Furthermore, a Kolmogorov–Smirnov test comparing the two distributions revealed a marginally significant difference between groups (*D* =.33, *p* =.055). These results suggest that the significant group difference in conformity tendency observed in the dot-counting task cannot be attributed solely to a uniform shift in the average level of conformity. Rather, it appears to be driven by a distinct subgroup (n = 9) of individuals with ASD who never changed their initial ratings in the dot-counting task (ASD-never-conforming), resulting in a skewed distribution. By contrast, the distribution of conformity tendencies observed in the preference rating task did not significantly deviate from normality in either group (ASD: *W* =.98, *p* =.840; NT: *W* =.97, *p* =.521), and there was no significant difference in distribution shape between groups (*D* =.10, *p* =.997; Figure 4A).

**Figure 4.**
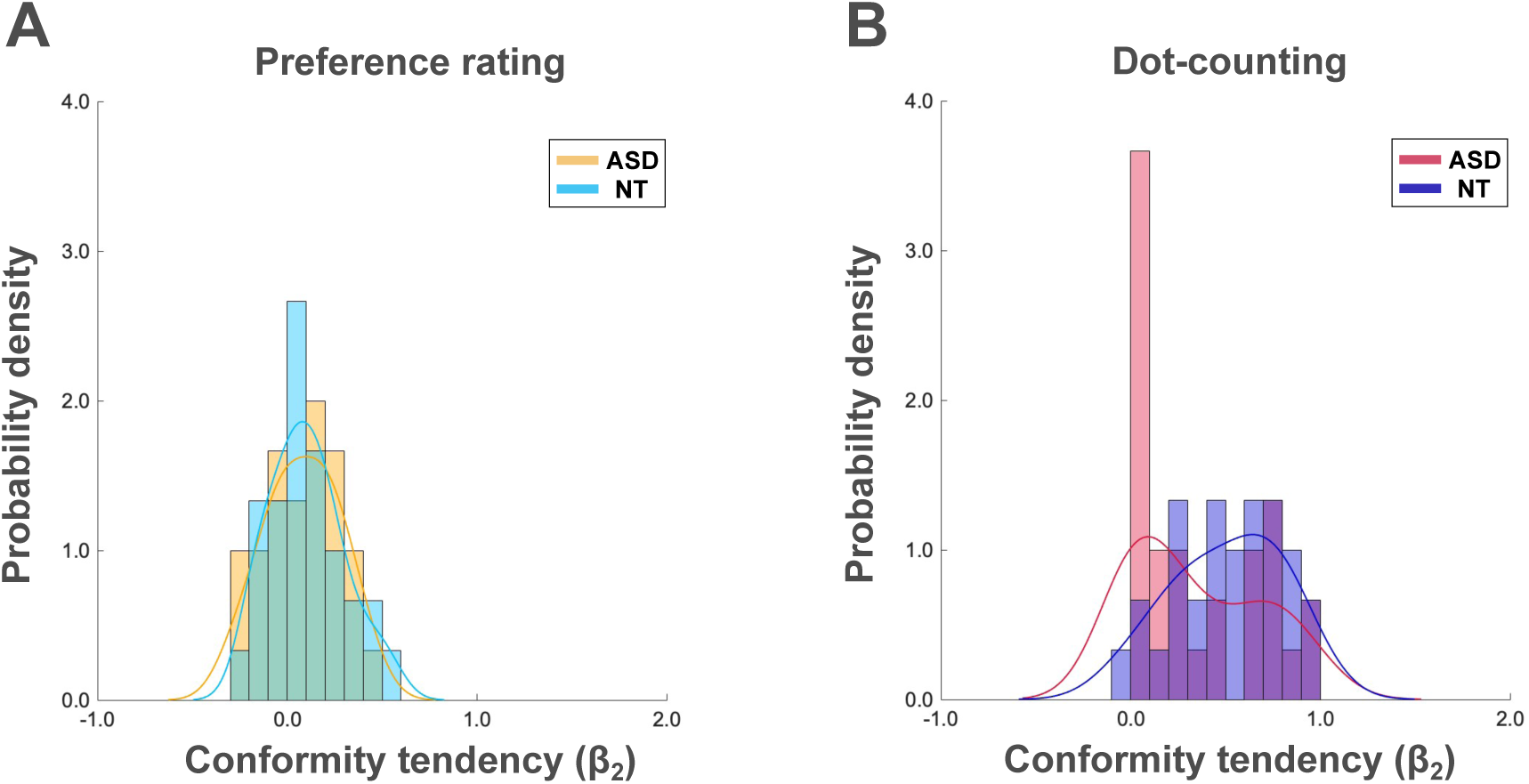
Probability distributions of conformity tendency Probability distributions of conformity tendency (β₂; derived from Eq. 1) are shown for each task using kernel density estimation (KDE; Gaussian kernel) and overlaid density-scaled histograms (bin width = 0.1). (A) Probability distribution of conformity levels in the preference rating task. The x-axis represents conformity tendency (β₂), and the y-axis denotes probability density (area = 1), reflecting the relative likelihood of observing a given level of conformity. (B) Probability distribution of conformity levels in the dot-counting task. The x-axis represents conformity tendency (β₂), while the y-axis denotes probability density (area = 1), reflecting the relative likelihood of observing a given level of conformity.

To statistically assess whether the original interaction effect (Figure 2 and Table 2) was driven by the ASD-never-conforming subgroup, we re-ran a group GLMM1 excluding the data from this subgroup. This analysis revealed a significant main effect of *Gap1O* (*b* = 0.46, *t* = 27.71, *p* <.001), indicating that greater discrepancy between self and group ratings continued to predict larger rating changes. However, the two-way interaction between *Gap1O* and *Group* was no longer significant (*b* = −0.01, *t* = −0.51, *p* =.609), nor was the three-way interaction between *Gap1O*, *Task*, and *Group* (*b* = 0.01, *t* = 0.16, *p* =.873). These results indicate that when the ASD-never-conforming subgroup is excluded, the ASD-conforming subgroup and NT controls exhibit comparable conformity tendencies across both tasks. This finding suggests that the original three-way interaction effect was primarily driven by the ASD-never-conforming subgroup.

It is possible that the complete lack of conformity in the dot-counting task observed among the ASD-never-conforming subgroup reflects an unusually strong consistency bias—that is, a rigid tendency to maintain one’s initial judgment—rather than a diminished sensitivity to social information per se. If this were the case, we would expect this subgroup to also exhibit high consistency in the preference rating task. To test this possibility, we calculated the correlation coefficients between initial and second ratings in the preference rating task for each participant, and compared these consistency bias indices across the ASD-never-conforming subgroup, the ASD-conforming subgroup, and the NT group using a one-way ANOVA. The analysis revealed no significant group differences (*F* (2, 57) = 1.05, *p* =.355, η² =.036), suggesting that consistency bias alone cannot account for the reduced conformity observed in the dot-counting task among the ASD-never-conforming subgroup.

## Discussion

In the current study, we employed two distinct tasks—the preference rating task and the dot-counting task—to investigate whether individuals with ASD exhibited different pattern of conformity based on distinct types of influence: normative and informational. We hypothesized that individuals with ASD would exhibit a reduced tendency to conform to others during the preference rating task, which involves subjective preference judgments and is primarily driven by normative influence, but would show a comparable level of conformity to NT individuals in the dot-counting task, where the presence of objectively correct answers, moderate task difficulty, and monetary incentives together make informational influence particularly relevant. Our results were the complete opposite of our predictions, the results revealed that while individuals with ASD exhibited a comparable tendency to conform during the preference rating task, their conformity was markedly diminished during the dot-counting task. Furthermore, we identified a subset of individuals with ASD who never modified their initial responses in the dot-counting task (ASD-never-conforming). That is, individuals in the ASD-never-conforming subgroup showed no conformity even in a context where social information was informative, explicit, and financially incentivized (dot-counting task task). Our data showed that this subgroup entirely explained the group differences in conformity tendency in the dot-counting task.

In the preference rating task, contrary to our hypothesis, individuals with ASD exhibited a conformity tendency comparable to that of NT individuals. This finding suggests that individuals with ASD shared similar underlying motivations for conformity as NT participants. However, an alternative explanation is that the motivation underlying conformity in ASD differs in nature. Rather than reflecting a genuine desire to align beliefs with others, conformity in ASD may represent an effort to appear socially competent and mask social difficulties—a phenomenon referred to as “social camouflaging” (Hull et al., 2017). Camouflaging has been described as a strategic adjustment of outward behavior to align with others while privately maintaining divergent attitudes, resembling “public conformity (or public compliance)” (Dittes & Kelley, 1956; Menzel, 1957). Unlike typical conformity processes that may culminate in private conformity (private acceptance)—a genuine shift in attitudes—camouflaging rarely entails true endorsement. From this perspective, the conformity tendencies observed in ASD during the preference rating task may have been driven by strategic self-presentation rather than actual belief change. Observing the normative cue (the displayed “average”) may have elicited a sense of social-evaluative risk, motivating individuals with ASD to modify their outward responses to appear “majority-consistent” and avoid being perceived as socially atypical or as having a developmental disorder, while privately maintaining their original opinions.

Thus, conformity in the preference rating task among individuals with ASD may be best characterized as public compliance achieved through social camouflaging, whereas for NT individuals, it likely reflects genuine private acceptance. Consistent with this interpretation for NT individuals, our previous work using the same t-shirt rating paradigm demonstrated that NT participants’ preferences remained influenced by others’ opinions even months later (Izuma & Adolphs, 2013), indicating private acceptance. Similarly, a neuroimaging study with NT individuals showed that others’ opinions modulated activity in reward-related brain regions, such as the ventral striatum—suggesting that social influence can alter the neural representation of subjective value (i.e., reflecting a genuine change in preference) (Zaki et al., 2011). Future studies should therefore test whether conformity in the preference rating task among individuals with ASD is accompanied by private acceptance (e.g., long-term preference change or changes in neural activity) or whether it remains limited to public compliance. Another promising research direction would be to examine whether conformity in ASD is associated with elevated stress, given that conforming may itself be psychologically burdensome when it functions as a strategy to mask autistic traits.

Importantly, with repeated use across the lifespan, camouflaging may become automatized and even operate unconsciously (Lawson, 2020). This possibility is particularly relevant to our paradigm: because the preference rating task contained a delay between the initial rating, the presentation of others’ ratings, and the subsequent rating, it is implausible that participants were consciously remembering and deliberately adjusting their responses. Thus, the conformity we observed in ASD may reflect automatic or internalized camouflaging processes, which outwardly resemble NT individuals’ conformity but are likely driven by different underlying motivations.

This camouflaging account may also explain the discrepancy between the present findings and those of Yafai et al. (2014), who examined conformity in children with ASD using a child-friendly adaptation of the classic Asch (1952) paradigm, where normative influence is salient. That study reported reduced conformity in children with ASD compared with controls, whereas in the present study with adults, we found no significant group difference in the preference rating task. One plausible explanation is developmental change: accumulated social experiences across the lifespan may shape compensatory strategies, such as camouflaging, that enable adults with ASD to adapt more effectively in socially evaluative contexts than children.

Unlike in the preference rating task, the dot-counting task revealed that individuals with ASD exhibited significantly reduced conformity compared with NT individuals. Closer inspection revealed that this group-level difference was not due to a uniform shift in central tendency but was driven by a distinct ASD subgroup of participants who never conformed (ASD-never-conforming). When this subgroup was excluded, the remaining participants with ASD (ASD-conforming subgroup) exhibited conformity tendencies comparable with NT individuals. These results suggest that the reduced tendency to conform under informational influence observed in the ASD group are not universally applicable to all individuals diagnosed with the condition, consistent with the broader recognition that ASD is highly heterogeneous, as demonstrated across phenotypic, neuroimaging, and etiological levels (e.g., Mottron & Bzdok, 2020). Prior work also points to such subgroups. Using the classic Asch (1952) paradigm, Bowler and Worley (1994) found that 50% of participants with ASD (4 out of 8) never conformed to confederates’ incorrect judgments. By contrast, this “no-conformity” pattern was observed in only 10% of typical control participants (1 out of 10) and in none of the verbal-IQ–matched control participants (0 out of 9). Although normative influence likely drove conformity in their paradigm and informational influence was more salient in ours, both tasks share two features that may help explain the emergence of a non-conforming subgroup: (i) both tasks involved perceptual decision-making; and (ii) conformity was explicit, so participants were aware when they did or did not conform—that is, when they chose to trust or disregard others’ estimates.

Building on the first point, our finding resonates with accumulating evidence that autistic populations are highly heterogeneous in perceptual style. Previous studies have identified subgroups with more pronounced difficulties in global integration or stronger reliance on perceptual detail (Robertson & Baron-Cohen, 2017; Simmons et al., 2009). The ASD-never-conforming subgroup in our study may thus represent individuals whose perceptual style is particularly resistant to incorporating socially derived information. Several theoretical accounts converge on this interpretation. From the perspective of weak central coherence (WCC; Happé & Frith, 2006), conformity in our task required participants to integrate their own perceptual estimates with others’ average response—a process of global information integration.

The diminished adjustment in the ASD group may therefore reflect a relative weakness in combining local and global sources of information into a coherent whole. The enhanced perceptual functioning (EPF) model (Mottron et al., 2006) offers a different emphasis, proposing that individuals with ASD show superior autonomy of low-level perceptual processes. On this account, the subgroup may have relied disproportionately on their own perceptually grounded estimates, making them less likely to revise judgments based on socially derived, abstracted information such as others’ average responses. Finally, the Bayesian account of autistic perception (Pellicano & Burr, 2012) suggests that individuals with ASD assign less weight to prior knowledge and depend more heavily on immediate sensory evidence. In this framework, others’ responses can be construed as socially derived priors that are given little weight. Taken together, despite their different emphases, these perspectives converge on the idea that the ASD-never-conforming subgroup prioritizes direct sensory evidence over socially derived information.

The second point—the explicit nature of conformity in our dot-counting task—may also be critical. Because participants were necessarily aware of whether they trusted or disregarded others’ estimates, subgroup differences may partly reflect variation in metacognitive processes. Specifically, the ASD-never-conforming subgroup may have misjudged the accuracy of their initial ratings, leading to unwarranted confidence in their own responses or skepticism about the reliability of others’ judgments. Prior research shows that higher confidence is generally associated with reduced susceptibility to informational influence (e.g., Fleming et al., 2018). In our dot-counting task, although individuals with ASD exhibited larger deviations from the actual number of dots than NT individuals, they nonetheless showed a reduced tendency to conform. This pattern suggests that the ASD-never-conforming subgroup may have placed excessive trust in their own estimates or discounted the reliability of others’ judgments, despite lacking objective grounds to verify either. Difficulties in evaluating one’s own performance, or reduced metacognitive accuracy, have been documented in ASD even when feedback is available (e.g., Grainger et al., 2014). McMahon et al. (2016), for example, used a face-emotion recognition task with trial-wise 1–5 confidence ratings (piecemeal facial information). Despite comparable accuracy to NT controls, the authors found that individuals with ASD exhibited both higher mean confidence and a wider confidence distribution, with more frequent use of extreme high-confidence categories—a “high-confidence/high-variance” profile rather than a uniform shift in mean confidence. This entails elevated confidence with weaker confidence–accuracy coupling (poorer metacognitive calibration). Taken together, these findings support heterogeneity in metacognition in ASD and are consistent with the presence of an overconfidence-biased subgroup, which may help explain non-conforming subgroups in our data.

Importantly, when the ASD-never-conforming subgroup was excluded, the remaining participants with ASD conformed to a similar extent as NT individuals, consistent with the findings of Lazzaro et al. (2019). Taken together, these observations imply that apparent group-level reductions in conformity may be better understood as reflecting heterogeneity within the ASD population. An important implication of the present findings is the need to identify distinct subgroups within the autistic population. Subgroup identification is increasingly recognized as essential for advancing understanding of etiology and underlying mechanisms, and for moving toward a personalized medicine approach in ASD research (e.g., Ousley & Cermak, 2014).

Because conformity is tightly linked to social adaptation, distinguishing between ASD-never-conforming and ASD-conforming groups may also inform targeted interventions and support strategies to enhance social participation. In this study, we did not have the data necessary to determine the factors that differentiate these subgroups, leaving the underlying mechanisms largely unexplored. Future research should seek to identify the factors—whether cognitive, motivational, or social—that give rise to these divergent conformity patterns.

A further, tentative possibility is that camouflaging may have contributed to some cases of conformity in the dot-counting task. Although anonymity was assured and the task did not include an explicit evaluative context, it is plausible that some participants nevertheless felt that their responses were noticed or could be judged (e.g., by the experimenter or an imagined audience). Because our experimental design allowed participants to revise their initial estimates explicitly and immediately, the ease of aligning with the displayed group average might have made impression management more likely for some participants. At the same time, informational motives cannot be ruled out. Even though no feedback about accuracy was provided, some participants may have assumed that the others’ average was more accurate and adjusted their responses accordingly. Thus, the conformity observed in the ASD-conforming subgroup during the dot-counting task may have been influenced by social camouflaging processes intended to manage social evaluation, although it could also, as in NT individuals, have arisen from a genuine motivation for accuracy. However, because camouflaging is typically motivated by concerns about acceptance and rejection, related processes may be more directly involved in normative contexts, such as with preference judgments (preference rating task), than in the accuracy-focused dot-counting task; accordingly, its influence in dot-counting is likely weaker. This distinction implies that camouflaging in ASD may manifest differentially depending on whether the context is socially evaluative or informational, and it specifies boundary conditions that future work should test.

One of the limitations of the current study is that our tasks involved structural differences. The preference rating task comprised two distinct sessions: others’ opinions were presented only in the first session, and participants were not informed in advance about the existence of the second session. Consequently, they were largely unable to make intentional decisions to conform to others. By contrast, in the dot-counting task, participants were asked to adjust their initial responses immediately after the presentation of others’ opinions, allowing them to intentionally make decisions to conform. These differences mean that the two tasks may not provide perfectly parallel tests of normative versus informational influence, and this structural asymmetry may partly contribute to the divergent behavioral patterns we observed. Further, the dot-counting task design may limit its suitability as a pure measure of informational influence. Participants estimated the number of presented dots using a sliding scale, and others’ opinions were displayed on the same scale during the task. Another limitation is that participants’ own and others’ responses were presented on the same response scale. This layout might invite adjustment based on visual distance rather than reliability-weighting (Öztel & Balcı, 2024). However, because the same format was consistently used in both tasks, such scale-related bias should be shared, and thus does not undermine the validity of our between-task comparison.

In conclusion, the current study identified that individuals with ASD exhibited different levels of susceptibility to distinct types of social influence: informational and normative. Whereas their conformity under normative influence appeared comparable to that of NT individuals—potentially reflecting camouflaging strategies—their conformity under informational influence was reduced, a pattern driven by a distinct subgroup who never modified their initial responses despite being provided with explicit and incentivized cues. The findings highlight the heterogeneity of conformity behavior in ASD and suggest that reduced conformity is not a universal trait but reflects variability in perceptual style, metacognitive processes, and camouflaging strategies. More broadly, the results underscore the importance of distinguishing between normative and informational influence in understanding social behavior in ASD. Future research should clarify the mechanisms that differentiate conforming from non-conforming subgroups, and examine how camouflaging operates across implicit and explicit contexts. Such work will advance theoretical models of social influence and provide a more nuanced understanding of the diverse ways in which autistic individuals navigate social information.

## Acknowledgments

We would like to thank Dr. Ryuichi Tamai for his support in constructing the experimental programs. Our sincere gratitude extends to all participants and their parents, and to all colleagues at our institutions, for their generous support and insightful discussions. This work was supported by JSPS KAKENHI Grant Number 19K24680.

