## Supporting Information for "Informational and Normative Influence on Conformity in Autism"

### Supplementary Note 1: Preregistered group comparison of across-trial conformity tendencies

To measure participants' conformity tendency across-trials in the dot-counting task, we also conducted a multiple regression analysis for every participant individually, as follows:

$$\text{Gap1D}\ (t + 1) = 1 + \beta_3 \times \text{Gap1O(t)}\ldots\ \text{(S1)}$$

In the equation (S1), *t* represents the number of trials, and *Gap1O*(t) denotes the difference between the participant's first rating and the ratings from other individuals (other people's answer – participant's first answer) at trial *t*. *Gap1D*(*t* + 1) represents the difference between the participant's first rating and actual number of dots (actual number of dots – first rating) at trial *t* + 1. Then, we compared across-trial conformity tendencies (i.e., $\beta_3$) between groups using a two-sample t-test. We also examined whether the across-trial conformity tendencies were different from 0 in each group using a one-sample t-test. We applied the conventional significance threshold $p < .05$.

### Supplementary Note 2: Group comparison of across-trial conformity tendencies using GLMM analysis

To conduct a group comparison on across-trial conformity tendencies in the dot-counting task, we also constructed a GLMM, as follows:

$$\text{Gap1D}\ (t + 1) \sim 1 + \text{Gap1O}(t) + \text{Gap1O}(t)\text{:Group} + (1 \mid \text{Participant})\ldots\text{(GLMMS1)}$$

where $t$ represents the number of trials, and *Gap1O(t)* denotes the difference between the participant's first rating and the ratings from other individuals (other people's answer – participant's first answer) at trial $t$. *Gap1D(t+1)* represents the difference between the participant's first rating and actual number of dots (actual number of dots – first rating) at trial $t + 1$.

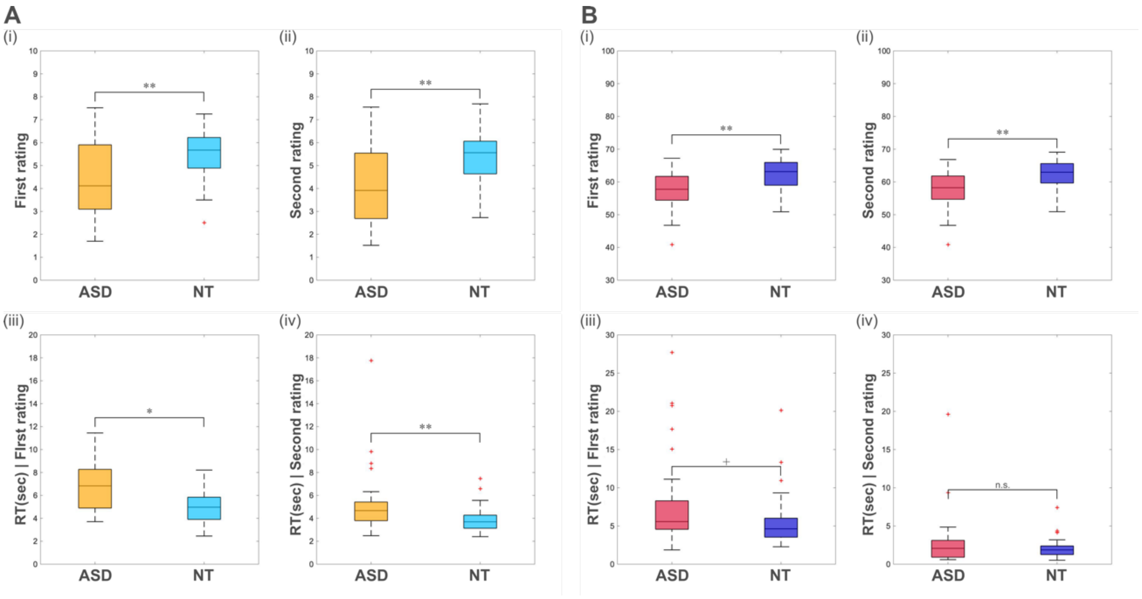


**Figure S1 Results of preregistered group comparison of measured variables**

A. Results of group comparison of measured variables in the preference rating task. (i) Participants' first ratings of the preference rating task are illustrated. Individuals with ASD displayed significantly lower preferences than NT individuals for the second ratings ($t$ (58) = −3.72, $p$ = .001). (ii) Participants' second ratings of the preference rating task are illustrated. Individuals with ASD reported significantly lower preferences than NT individuals for the second ratings ($t$ (58) = −3.85, $p$ = .001). (iii) Response times for the first ratings of the preference rating task are illustrated. Individuals with ASD responded significantly slower than NT individuals for the first rating ($t$ (58) = 2.62, $p$ = .014). (iv) Response times for the second ratings of the preference rating task are illustrated. Individuals with ASD responded significantly slower than NT individuals for the first rating ($t$ (58) = 3.21, $p$ = .003). B. Results of group comparison of measured variables in the dot-counting task. (i) Participants' first ratings of the dot-counting task are illustrated. Individuals with ASD estimated the number of dots to be lower than NT individuals for the first ratings ($t$ (58) = −2.97, $p$ = .006). (ii) Participants' second ratings of the dot-counting task are illustrated. Individuals with ASD estimated the number of dots to be significantly lower than NT individuals for the second ratings ($t$ (58) = −2.86, $p$ = 0.008). (iii) Response times for the first ratings of the dot-counting task are illustrated. Individuals with ASD took a marginally significantly longer time than NT individuals ($t$ (58) = 1.94, $p$ = .062). (iv) Response times for the second ratings of the dot-counting task are illustrated. Individuals with ASD determined their final answer (whether to maintain or revise their first response) as quickly as NT individuals ($t$ (58) = 1.26, $p$ = .219). The box indicates the interquartile range (25th–75th percentile), and the central line represents the median. Whiskers extend to 1.5 × IQR, and red crosses indicate outliers. $^{**}p < .001$, $^{*}p < .05$, $+\,p < 0.1$, n.s.: not significant.

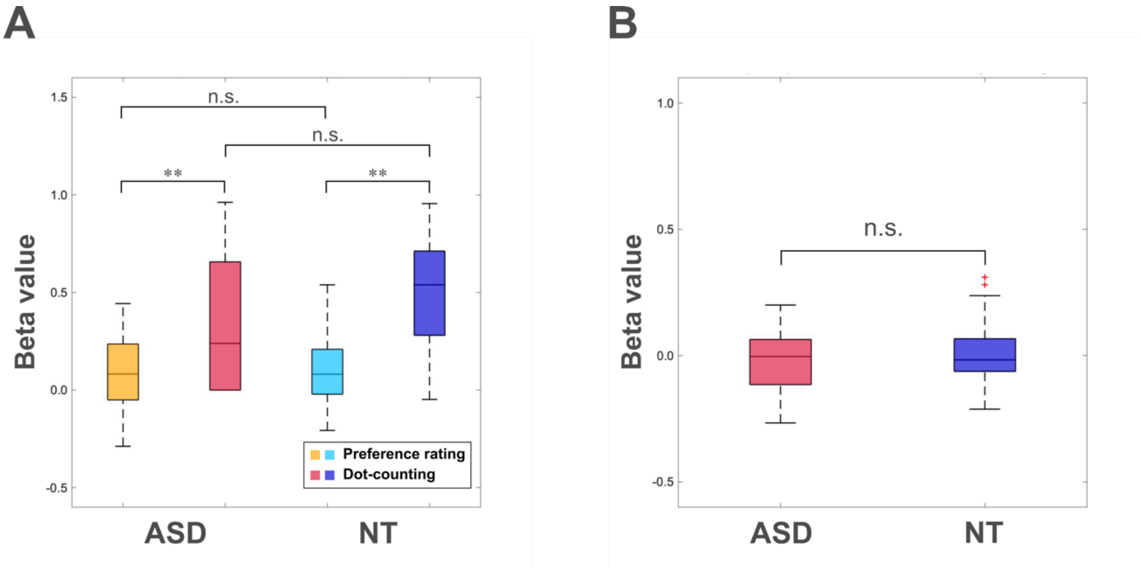


**Figure S2 Preregistered group comparison of conformity tendencies**

Results of the preregistered group comparison of conformity tendencies. (A) Results of the mixed two-way ANOVA on conformity tendencies. The conformity tendencies (calculated as $\beta_2$ from Equation 1) were significantly greater than 0 in both groups across both tasks (preference rating: ASD: $t$ (29) = 2.29, $p$ = .029; NT: $t$ (29) = 2.85, $p$ = .008; dot-counting: ASD: $t$ (29) = 5.54, $p$ < .001; NT: $t$ (29) = 9.79, $p$ < .001). Subsequently, we compared participants' conformity tendencies ($\beta_2$) using a mixed two-way ANOVA, with group (ASD and NT) as a between subject factor and task type (preference rating and dot-counting task) as a within-subject factor. As a result, we found marginally significant main effects of Group ($F$ (1,58) = 3.23, $p$ = .077, $\eta_p^2$ = .053) and Task ($F$ (1,58) = 60.32, $p$ < .001, $\eta_p^2$ = .51). The Group by Task interaction trended toward significance ($F$ (1,58) = 2.89, $p$ = .094, $\eta_p^2$ = .047). The simple main effects analysis further demonstrated that Task had a significant effect in both groups, indicating that conformity tendency in the dot-counting task was stronger than in the preference rating task in both groups (ASD: $F$ (1,58) = 18.40, $p_{adjusted}$ < .001, $\eta_p^2$ = .241; NT: $F$ (1,58) = 44.81, $p_{\text{adjusted}}$ < .001, $\eta_p^2$ = .436). However, the simple main effect of Group was not significant in either the preference rating or dot-counting task (preference rating : $F$ (1,58) = 0.15, $p_{adjusted}$ = 1.000, $\eta_p^2$ =.002; dot-counting : $F$ (1,58) = 4.22, $p_{\text{adjusted}}$ = .178, $\eta_p^2$ = .068). (B) Results of the group comparison of across-trial conformity tendencies in the dot-counting task. The across-trial conformity tendencies (calculated as $\beta_3$ from Equation (S2)) in the dot-counting task were not different from 0 in both groups (ASD: $t$ (29) = −0.69, $p$ = .498; NT: $t$ (29) = 0.49, $p$ = .625). We did not find significant group difference ($t$ (58) = -0.75, $p$ = .460). The box indicates the interquartile range (25th–75th percentile), and the central line represents the median. Whiskers extend to 1.5 × IQR, and red crosses indicate outliers. **$p$ < .001, n.s.: not significant.

**Table S1 Results of GLMMS1 examining group differences in across-trial conformity tendencies**

| | Estimate | SE | $t$ | $p$ | 95% CI |
|---|---|---|---|---|---|
| Intercept | 0.00 | 0.01 | 0.00 | 1.000 | [−0.03 0.03] |
| Gap1O | 0.01 | 0.02 | 0.67 | .503 | [−0.02 0.05] |
| Gap1O:Group | −0.03 | 0.03 | -1.01 | .311 | [−0.08 0.02] |

*Note.* Gap1O: difference between first rating and others' rating, Group: dummy code of group (ASD:1; NT:0), SE: standard error, CI: confidence interval.
